# Highly Parallel Tissue Grafting for Combinatorial In Vivo Screening

**DOI:** 10.1101/2023.03.16.533029

**Authors:** Colleen E. O’Connor, Anna Neufeld, Chelsea L. Fortin, Fredrik Johansson, Jonathan Mene, Sarah H. Saxton, Susana P. Simmonds, Irina Kopyeva, Nicole E. Gregorio, Cole A. DeForest, Daniela M. Witten, Kelly R. Stevens

## Abstract

Material- and cell-based technologies such as engineered tissues hold great promise as human therapies. Yet, the development of many of these technologies becomes stalled at the stage of pre-clinical animal studies due to the tedious and low-throughput nature of *in vivo* implantation experiments. We introduce a ‘plug and play’ *in vivo* screening array platform called Highly Parallel Tissue Grafting (HPTG). HPTG enables parallelized *in vivo* screening of 43 three-dimensional microtissues within a single 3D printed device. Using HPTG, we screen microtissue formations with varying cellular and material components and identify formulations that support vascular self-assembly, integration and tissue function. Our studies highlight the importance of combinatorial studies that vary cellular and material formulation variables concomitantly, by revealing that inclusion of stromal cells can “rescue” vascular self-assembly in manner that is material-dependent. HPTG provides a route for accelerating pre-clinical progress for diverse medical applications including tissue therapy, cancer biomedicine, and regenerative medicine.

## Main

Material- and cell-based technologies designed for therapeutic transplantation hold enormous potential for treating diverse human diseases. Over the past several decades, a vast portfolio of such technologies has been developed^1^, including materials designed to support angiogenesis^2–4^, promote transplanted cell survival^5,6^ or fate choice^7,8^, modulate the immune response^9,10^, capture malignant cells^11^, or deliver biological therapeutics^12,13^. More recently, numerous technologies composed of materials and cells, such as engineered tissues designed to replace or supplement the functions of highly metabolic tissues and organs, have been introduced^14^. Yet, most of these technologies languish in pre-clinical stages of development^15,16^. During the pre-clinical stage, the biggest bottleneck is arguably at the stage of *in vivo* studies, when technologies are surgically implanted into an animal (host) to assess their safety and efficacy.

This bottleneck is particularly formidable for technologies that combine both material- and cellular-elements, such as engineered tissues and living materials. In these fields, advances at the interfaces of material science, additive manufacturing, and cell biology have led to massive progress, such as towards fabricating 3D printed human organs and self-assembly of organoids, or “mini-organs”, in a dish^17–21^. In such technologies that are intended to be surgically implanted, each of the material and cellular components has the potential to interact with the biology of the host upon implantation^1^. That is, each component has the potential to contribute to an implant’s biological compatibility, degree of integration with host tissue such as vascular connectivity, support of transplanted cell viability and assembly, and other factors critical to the implant’s safety and efficacy^1,22^. Some emerging implantable technologies go so far as to hijack or leverage aspects of host biology to enhance a technology’s therapeutic outcome^23–29^. Thus, animal models remain a particularly steep requirement for combination technologies aiming to progress from bench to bedside.

Despite the critical role of the host in implant outcome for each of these technologies, scientific understanding of host biology remains incomplete. This knowledge gap, or “black box”, means that the formulations of material- and cell-based implants must still be tested and optimized empirically *in vivo.* The sheer number of material and cellular permutations possible, together with the inherent complexity and low-throughput nature of animal experiments, in which a single implant is typically surgically implanted into a given living host, has meant that progress slows to a crawl at the stage of pre-clinical animal studies (Fig. 1a). Thus, evaluating all possible formulations *in vivo* often remains experimentally intractable. New methods that overcome the time, financial capital, throughput, and ethical issues that arise in pre-clinical surgical implantation studies would massively accelerate regenerative medicine^30–32^.

**Figure 1.**
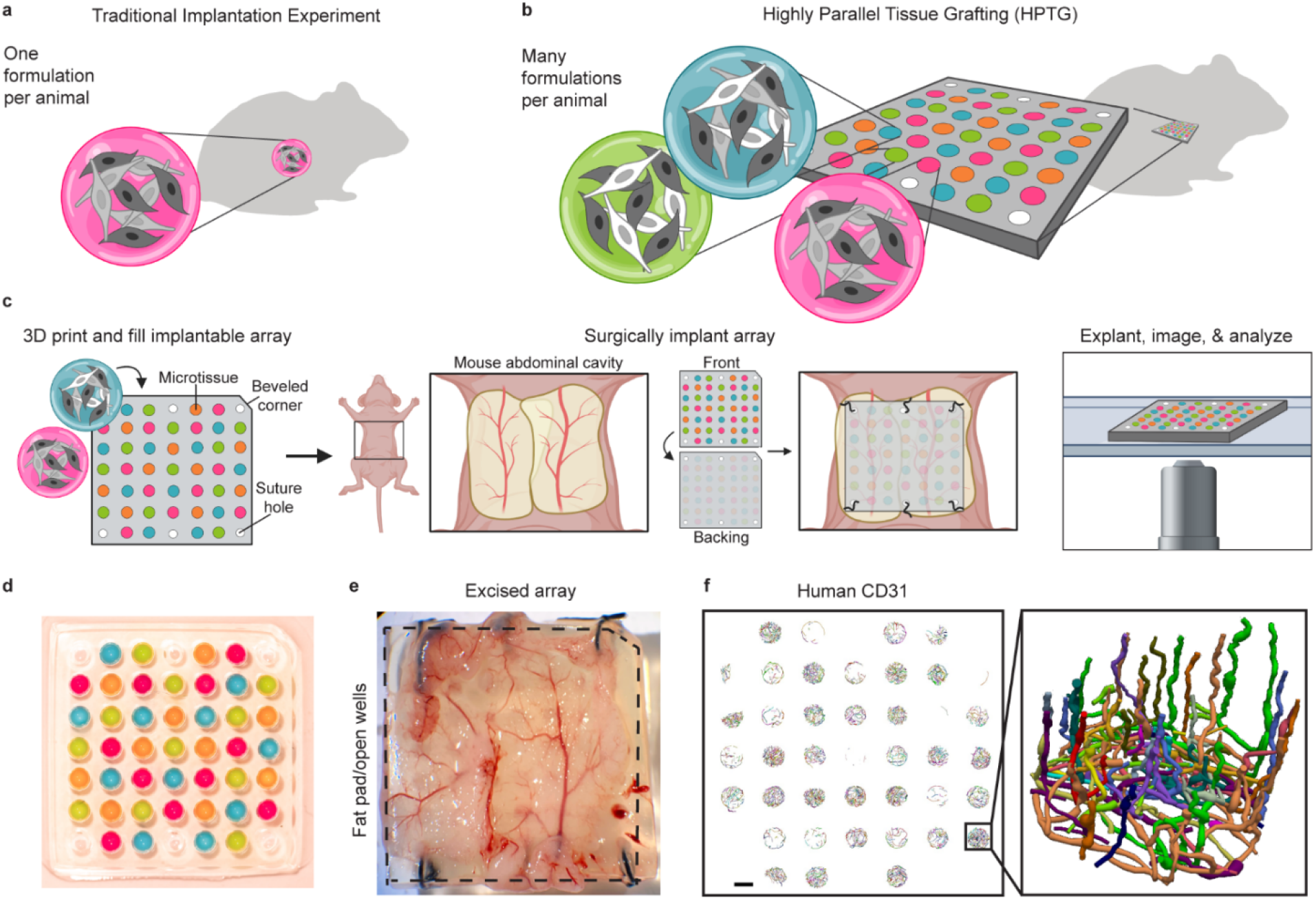
Development of Highly Parallel Tissue Grafting for in vivo screening. **a,** Schematic of a traditional *in vivo* experiment where only one formulation is typically tested in a given animal host. **b,** Schematic of HPTG platform, in which many material and/or cellular formulations are screened in parallel within a single animal host. **c,** Schematic of HPTG workflow (left to right) depicting 3D printing and filling HPTG array (left), surgically implanting array into abdominal space of mouse host (center), and then explanting, imaging, and analyzing array in 3D (right). **d,** A photograph (gross image) of a 3D printed array, in which wells in the array are filled with colored dyes. **e,** A gross image of an HPTG array explanted after 1 week *in vivo*, demonstrating the host fat pad adhered across the top of the open wells of the array. **f,** (Left) HPTG array with 3D reconstructed human (huCD31+) self-assembled networks within each well. Scale bar, 1000 μm. (Right) Angled orientation of a single microwell showing a 3D reconstructed network (huCD31). Individual colors represent individually segmented microvessels. The angled view indicates human endothelial vessels protruding up and away from the microwell bottom.

At the stage of *in vitro* studies, previous revolutions in miniaturizing biological assays, such as advancements in multiwell plates, robotic liquid handling, microarray technology, and microfabrication ushered in a new era of parallelized and combinatorial screening. This progress enabled higher throughput *in vitro* studies and selection of biological entities such as small molecules, DNA, extracellular matrices, biomaterials, and microenvironments^33–38^. We postulate that analogous approaches for screening material- and cell-based technologies at the *in vivo* stages of product development would accelerate pre-clinical progress.

Here, we developed Highly Parallel Tissue Grafting (HPTG), a new method for interrogating designer material- and cell-based environments *in vivo* at scale (Fig. 1b). This method adapts 3D printing technology to fabricate implantable arrays with individually addressable material- and cell-based environments (Fig. 1b). We use HPTG to uncover how the combinatorial material- and cell-based environments govern vascular network self-assembly and vascular graft-host inosculation *in vivo.*

### Development of highly parallel tissue grafting platform

We recently developed a stereolithography apparatus for tissue engineering (SLATE), which enabled 3D printing of biocompatible hydrogels to create complex topologies volumetrically by converting photoactive liquids into structured parts through localized photopolymerization^17^. We identified photopolymerizable hydrogel formulations that can be 3D printed using SLATE to create engineered tissues, which could be successfully implanted in animals for several weeks^17^. While this and earlier work in the 3D bioprinting field has focused on fabricating therapeutic engineered tissues and organs (Fig. 1a), we reasoned that this 3D printing technology could be adapted to fabricate biocompatible devices for *in vivo* screening (Fig. 1b).

Towards this end, we fabricated a 3D printed HPTG device with individually addressable wells for parallelized and combinatorial screening that can be implanted *in vivo* (Fig. 1b-d). In this device, each individually addressable well can be filled with different material- and cell-based formulations, which can be retained in the device after implantation and for downstream analyses, such as biomedical imaging (Fig. 1c). Formulations in each well are comprised of any user-defined combination of materials (*e.g*., hydrogels, extracellular matrices), cellular composition (e.g., cell type(s), density, ratio), or biological molecules (Fig. 1c). The studies here will use both material and cellular components, thus we henceforth will call the formulations within each well of the HPTG device “microtissues” (Fig. 1b,c).

The HPTG device is composed of water, gelatin methacrylate [GelMA, 10 weight % (wt%)], and poly(ethylene glycol) diacrylate [PEGDA, 3.4 kDa, 3.25 wt%], a biocompatible photopolymerizable hydrogel chosen such that the device will retain its structural integrity (i.e., not break or crack) and remain pliable to facilitate direct apposition of microtissue formulations with host tissue, even with host movement after surgical implantation. Inclusion of PEGDA imparts mechanical strength to the hydrogel and inhibits degradation^39^, and inclusion of GelMA facilitates host apposition and subsequent downstream analysis techniques such as tissue sectioning.

We first leveraged the rapid prototyping capabilities of SLATE to fabricate and investigate design candidates suitable for *in vivo* screening arrays (Extended Data Fig. 1a). We created HPTG device designs using computer-aided design (CAD) and then 3D printed the devices using SLATE^17^. We first designed and tested a device architecture which contained an array of cylindrical wells that were open on both faces, such that when wells were filled, the interaction between microtissues and host tissue could occur on both sides of the device upon its implantation (Extended Data Fig. 1a, left). Once printed, the hydrogel array was washed for three days to swell to equilibrium and remove any unreacted monomer. We filled every well of the array with a fibrin hydrogel. The filled array was surgically implanted on perigonadal adipose tissue in the abdominal space of an athymic mouse, an implant site commonly used for material- and cell-based technology implantations due to its support of tissue vascularization and function^23,40–42^. Upon explant after one week, we found that host tissues (*i.e*., intestine) frequently engrafted and protruded through the wells in this open-well system, dislodging microtissues from the array (Extended Data Fig. 1b) and thus prohibiting downstream analysis on microtissues.

To overcome this obstacle, we added a backing component to the array architecture, which essentially created a hydrogel “well plate”, wherein each cylindrical well comprises one open face (Extended Data Fig. 1a, right). We postulated that upon implantation, this design would enable microtissues to directly interface with host tissue on one side, while the closed side prevents host tissue penetration through the array. To test the backing concept, we 3D printed devices with both designs: 1) HPTG arrays with open wells, and 2) HPTG arrays with backing on one side of the array (Fig. 1a-d). We then filled wells of these arrays with HEK 293T cells engineered with a constitutive luciferase reporter (for cell visualization downstream). Corner and center wells at the array edges were left unfilled to facilitate suturing. Arrays were then surgically implanted onto the perigonadal fat pad of mice. For the arrays with backing, the open side of the wells was faced down and thus directly apposed to the fat pad at the time of implantation (Fig. 1c, center). When the arrays were explanted after 1 week, we observed that explanted arrays with open wells (without the backing component) recovered minimal microtissue within the wells, similar to our previous studies (Extended Data Fig. 1b,c). Conversely, we observed full recovery of microtissues within the array wells with the backing (Extended Data Fig. 1c). We also observed host adipose tissue engrafted over the top of the wells, indicating capability of host tissue to interface directly with microtissue in the wells.

Having identified a well architecture that supports graft-host interactions and maintains device and microwell integrity *in vivo*, we next scaled the HPTG array design to increase the number of wells per array (Extended Data Fig. 1d). We ultimately settled upon an HPTG array configured by a grid of 7 by 7 wells, comprised of 6 suture holes and 43 individual microtissue wells and dimensions of 18 mm x 18 mm x 2 mm (Fig. 1c,d). The top-right corner of the device has a beveled edge that serves to define device orientation, and wells at the corners and center of array edges are left without backing and reserved for suturing (Fig. 1c,d). Each array is 3D printed in minutes and microtissue wells are then filled via standard pipetting (Fig. 1d). We confirmed that this array design supports robust engraftment, with direct apposition of the perigonadal fat pad to the array after explant of the HPTG array from the mouse host (Fig. 1e). Each well within the explanted arrays can then be visualized and computationally analyzed^43^ (Fig. 1c,f). This HPTG design thus opens the possibility of testing up to 43 different material- and cell-based microtissue formulations in a single mouse.

### Screening material microenvironments for vascular assembly in vivo

While major advances in tissue engineering have brought the field closer to producing complex highly metabolic solid tissues and organs for therapeutic transplant, tissue vascularization and integration with host circulation remain key barriers to their clinical translation^17,23,44–46^. One strategy for vascularizing engineered tissues relies on the inherent capacity of human endothelial cells seeded within a biomaterial to undergo morphogenesis. Specifically, several studies have shown that in some microenvironmental settings, endothelial cells can self-assemble to create vascular networks lined with both graft and host endothelial cells, connected to host vasculature, and carrying host blood^27,47,48^. We hypothesized that the capability of endothelial cells to undergo vascular self-assembly *in vivo* depends upon the formulation of the material microenvironment within which they are embedded^29,49–52^.

To test this hypothesis and apply HPTG, we profiled material environments for their ability to support vascular self-assembly *in vivo* (Fig. 2a). We focused our studies on GelMA, due to its biocompatibility^53^, bioactive moieties that support cell attachment^54^, and photoactive moieties that facilitate its usage with numerous biofabrication strategies, including 3D printing^55^. We suspended endothelial cells (primary Human Umbilical Vein Endothelial Cells, HUVECs) and stromal cells (primary Normal Human Dermal Fibroblasts, NHDFs) within 3 wt%, 5 wt%, 10 wt%, or 15 wt% GelMA microenvironments, which represent a range of stiffness (Extended Data Fig. 2a). We then dispensed these cellularized pre-polymers into wells to form microtissues, in a randomized manner across the array (Fig. 2a). After filling, the entire HPTG array was exposed to near-ultraviolet (UV, 405 nm) light to polymerize the cellularized GelMA microenvironments within the array wells^56,57^. After filling and polymerization, most cells within all GelMA microenvironments remained viable (Extended Data Fig. 2b).

**Figure 2.**
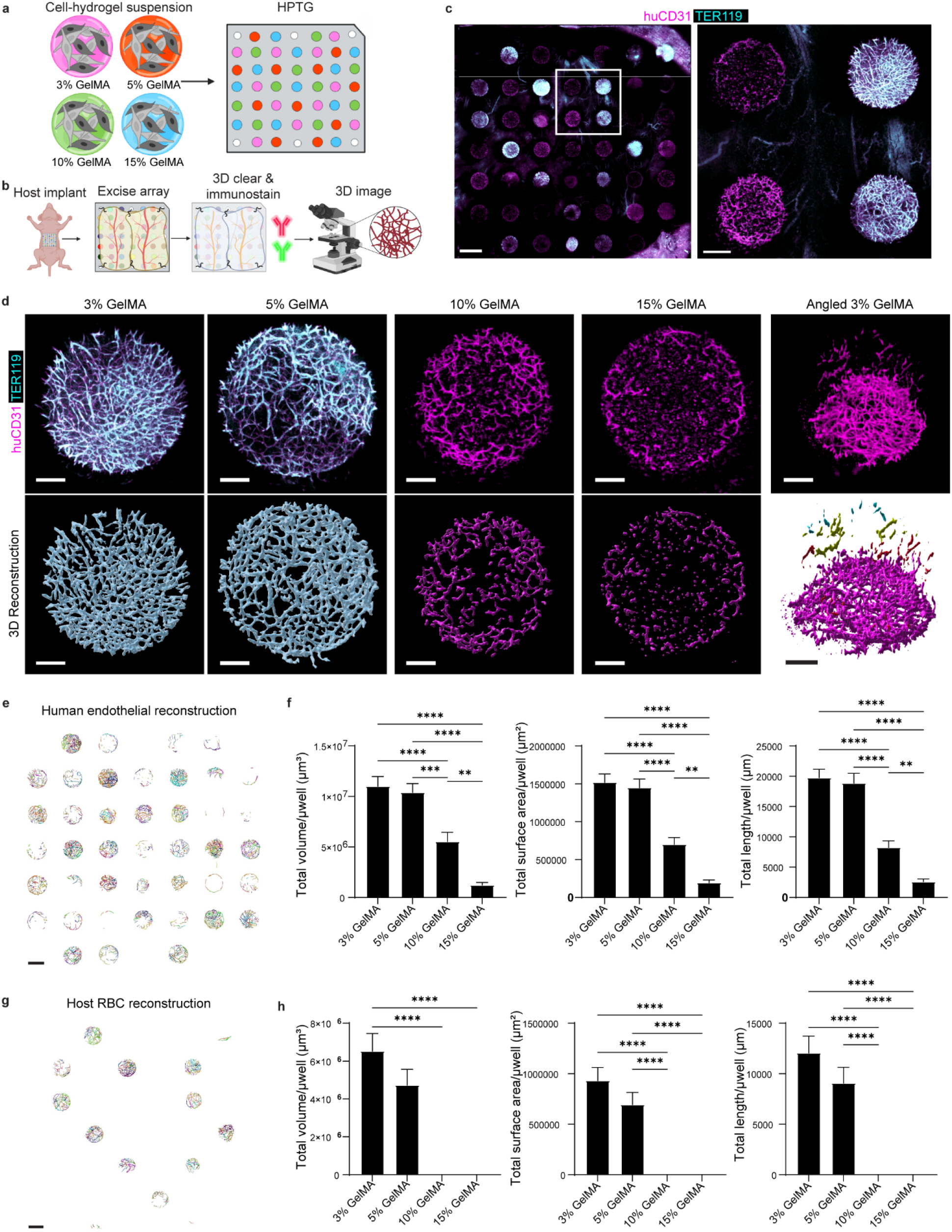
HPTG identifies material microenvironments that support vascular assembly in vivo. **a,** Schematic of experimental design, which assessed four different material formulations. In each HPTG device there were four materials screened, each with 11 technical replicates. There were n=4 mouse hosts, with each mouse receiving the same array layout. **b,** Schematic of post-implant workflow. HPTG devices are excised from the mouse host, 3D cleared and immunostained, and then 3D imaged using confocal microscopy. **c,** (Left) Max intensity projection (MIP) of an explanted HPTG device immunostained with huCD31 (human endothelial cells; magenta) and TER119 (red blood cells; cyan) and cleared array explanted after 1 week in vivo. Scale bar, 1500 μm. (Right) MIP magnification inset of four microwells. Scale bar, 400 μm. **d,** MIPs of representative microtissues stained for huCD31 (magenta) and mouse TER119 (cyan) representing the four microenvironments screened: 3 wt% GelMA, 5 wt% GelMA, 10 wt% GelMA, and 15 wt% GelMA. Scale bars, 200 μm. **e,** Image of reconstructed 3D huCD31+ microvascular self-assembled networks across an explanted HPTG array using Vesselucida 360 software. Scale bar, 1000 μm. **f,** Quantitative graphs of human (huCD31+) self-assembled network volume, surface area, and total length per microwell for each microenvironment screened. **g,** Image of reconstructed 3D mouse TER119+ microvascular self-assembled networks across an explanted HPTG using Vesselucida 360 software. Scale bar, 1000 μm. **h,** Quantitative graphs of mouse red blood cell+ self-assembled network (TER119+) volume, surface area, and total length per microwell for each microenvironment screened. Error bars indicate SEM. *P < 0.05 by one-way analysis of variance (ANOVA) followed by Tukey’s post-hoc test.

To investigate the ability of each GelMA microenvironment to support vascular self-assembly *in vivo*, we surgically implanted the HPTG arrays onto the perigonadal fat pad of athymic mice (Fig. 2b). After one week *in vivo*, the arrays were excised and cleared to render the HPTG array transparent and optically clear, to permit deeper imaging^58^. Cleared arrays were then 3D immunostained for human endothelial cells (huCD31, magenta) and mouse red blood cells (TER119, cyan) (Fig. 2b,c). Subsequent 3D image analysis of immunostained arrays revealed a range of self-assembled human endothelial networks within the various wells (Fig. 2c,d), with no evidence of vessel penetration between the walls of the wells (Fig. 2c,d).

To quantify vascular self-assembly in each material condition, we leveraged vessel visualization reconstruction software to parameterize huCD31+ and TER119+ networks across arrays (Fig.1f, 2e-g). Interestingly, we identified an inverse and dose-dependent relationship between endothelial cell network assembly and GelMA material concentration (Fig. 2f), with a 13-fold difference in assembled network volume between the lowest and highest wt% GelMA conditions (Fig 2d-f). Furthermore, 3 wt% and 5 wt% GelMA microenvironments supported infiltration of mouse red blood cells within human vascular microstructures, suggesting the inosculation of these networks with the host vasculature, whereas 10 wt% and 15 wt% GelMA microenvironments were mostly void of RBCs (Fig. 2d,g,h). These results were validated with 2D histological analyses, which demonstrated that 3 wt% and 5 wt% GelMA microenvironments contained microvascular structures filled with red blood cells, 10 wt% microenvironments had some sparse, hollow-lumened microvascular structures with no blood, and 15 wt% microenvironments contained only punctate cells, again with no blood (Extended Data Fig. 3). HPTG thus demonstrated that human vascular network self-assembly and inosculation with the host *in vivo* heavily depends upon formulation of the microenvironment that encases the grafted endothelial cells.

### An HPTG device design that overcomes potential position effects

While our initial study (Fig. 2) demonstrated the utility of HPTG, we next sought to increase the impact of HPTG for parallelized screening by increasing the number of conditions tested per animal. This required careful HPTG device design in order to overcome possible confounding due to spatial variability in the host, as well as attention to the sample size required to obtain adequate statistical power; we therefore next worked to address these two topics.

Because the array has a fixed number of 43 wells, increasing the number of conditions tested per animal leads to a decrease in the number of technical replicates per condition. Thus, testing more conditions per animal exacerbates the possibility that the technical replicates corresponding to a particular condition might be located in a small area of the array, leading to results that are confounded by a so-called “position effect”. That is, spatial variability of host features in the animal tissue (*e.g*., branching large vessels) or variable host tissue apposition across an array could affect the experimental outcome, and thus introduce bias in our results. Thus, we set out to further develop the HPTG platform to protect against this so-called “position effect”.

To do this, we designed a set of array layouts to ensure that every experimental condition appears in a variety of well positions across the study (Extended Data Fig. 4). This is achieved by creating different array layouts for each mouse in a given study. Once the first array has been designed, each subsequent array shifts the design of the first array by a random number of rows and columns (Extended Data Fig. 4a).

Concomitantly, we performed power analyses to determine how many biological and technical replicates would be needed to achieve high statistical power to detect effect sizes comparable to those seen in our initial vascular assembly data (Fig. 2). Our analyses showed that we could screen up to 21 conditions, with two technical replicates for each condition in a given array, using as few as eight mice (Extended Data Fig. 4b) and still detect effect sizes comparable to those seen within data in Fig. 2. Based on these analyses we created eight HPTG device maps, in which each array tests 21 experimental conditions with 2-3 technical replicates per array (Extended Data Fig. 4). We then set out to leverage this array design experimentally (Fig. 3).

**Figure 3.**
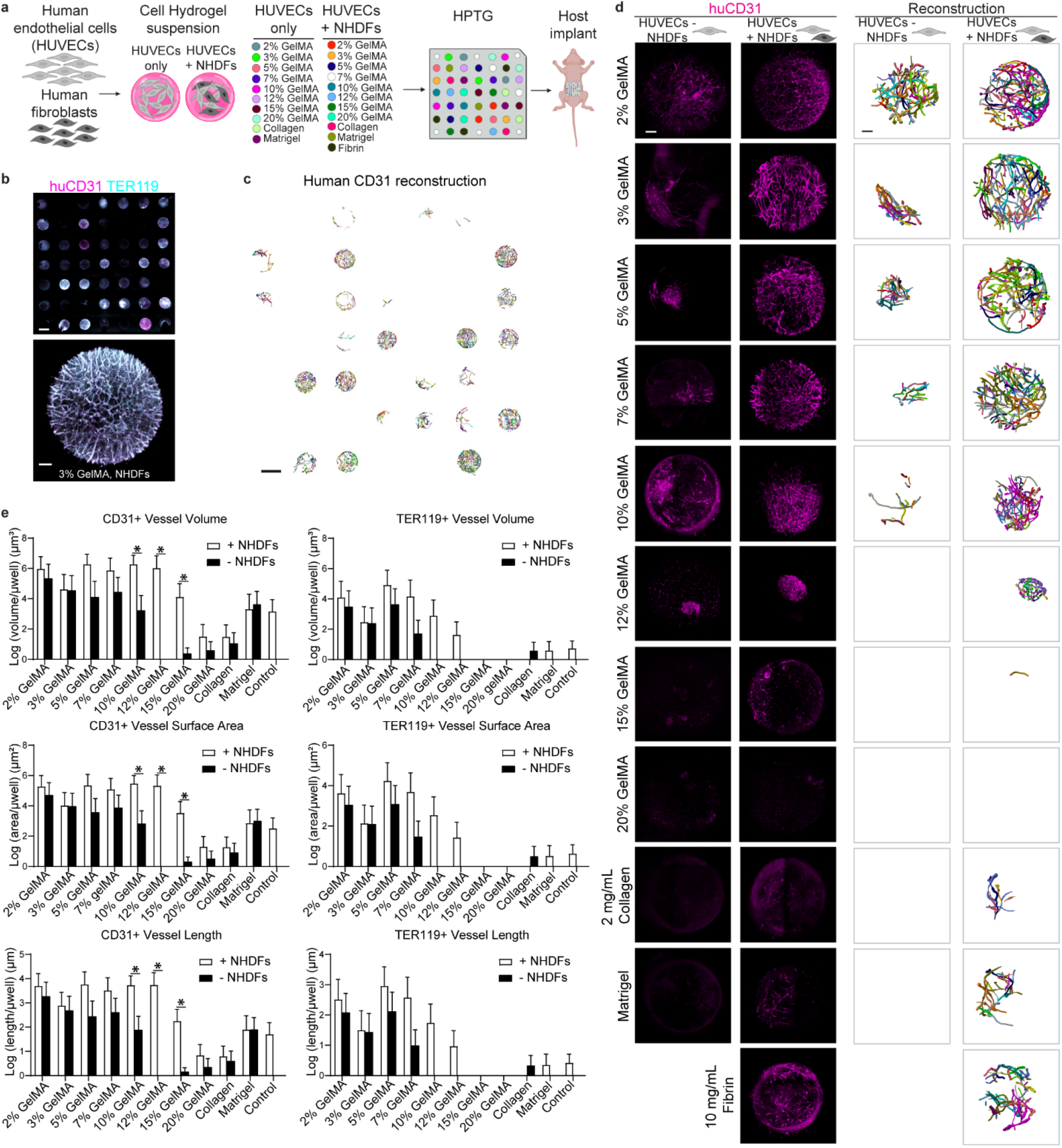
HPTG for combinatorial screening of cellular and material microenvironments in vivo. **a,** Schematic of experimental design, which screened 21 different material and cellular formulations, with 2-3 technical replicates per condition in each array. There were n=8 mouse hosts, with each mouse receiving a different array layout. **b,** Max intensity projection (MIP) of (top) a representative HPTG array from this experiment explanted after two weeks *in vivo*, immunostained for huCD31 (human endothelial cells; magenta) and TER119 (mouse red blood cells; cyan) and cleared. Scale bar, 1000 μm. (Bottom) MIP zoom in of a well containing HUVECs with NHDFs in a 3 wt% GelMA microenvironment. Scale bar, 100 μm. **c,** Image of reconstructed 3D huCD31+ microvascular self-assembled networks across an explanted HPTG array using Vesselucida 360 software. Scale bar, 1000 μm. **d,** (Left) MIPs of representative microtissues stained for huCD31 (magenta) within the 21 microenvironments screened. Scale bar, 100 μm. (Right) Corresponding huCD31+ self-assembled networks reconstructed in 3D using Vesselucida 360 software. Scale bar, 100 μm. **e,** Quantitative graphs of (left) human CD31+ and (right) TER119+ self-assembled network volume, surface area, and total length per microwell for each microenvironment screened. Error bars indicate SEM. *P < 0.05 after a Bonferroni correction for the difference between the coefficients for each given material condition +/- NHDFs, in a linear model with a fixed effect for each condition and each mouse.

### Scaling HPTG for combinatorial material and cellular screening in vivo

While most prior *in vivo* experiments have evaluated the influence of one variable at a time in implantable engineered tissues (*e.g*., material or cell formulation), microenvironmental ecosystems are composed of numerous interacting components, including extracellular matrices, cell populations such as stromal cells, and cell-secreted factors. We reasoned that the ability to screen many conditions *in vivo* would uniquely enable identification of the combinatorial effects of different classes of microenvironmental variables.

Thus, we next examined how combinatorial material-cellular interactions affect vascular assembly *in vivo*. Towards this end, stromal cells have been shown to enhance vascular self-assembly in engineered tissues upon their implantation in some settings^29,51,59^. These findings, coupled with our observation of the importance of material environment in vascular assembly (Fig. 2), made us seek to unveil the combinatorial effect of different material formulations and inclusion of stromal cells (e.g., fibroblasts) on vascular assembly *in vivo*. We encapsulated endothelial cells in the presence or absence of human fibroblasts in various materials, such that individual wells were filled with microtissues consisting of 21 different material and cellular combinatorial formations (Fig. 3a). These cells were encapsulated in an expanded portfolio of GelMA matrices, as well as a subset of natural hydrogels, including fibrin and collagen, two matrix components previously shown to support tissue vascularization^19,60,61^ and cell transplantation^62–65^ (Fig. 3a). Each experimental condition had 2-3 technical replicates within each array and arrays were laid out as denoted in Extended Data Fig. 4 to maximize power and minimize position bias.

Seeded HPTG arrays were implanted into mice for two weeks to test the impact of material and cell formulation on the formation and maintenance of self-assembled vascular networks lined with human endothelial cells *in vivo* (Fig. 3). After explant of the HPTG arrays at two weeks, we immunostained and volumetrically imaged the arrays to visualize 3D human endothelial cell self-assembly (huCD31) and host red blood cells (TER119). We then computationally reconstructed and parameterized networks across the arrays from each mouse (Fig. 3b-d).

As mentioned earlier, we used a different array layout in each mouse (Fig.3, Extended Data Fig. 4), to protect against potential confounding by “position effect”, i.e., the possibility that the position of a given well (microtissue) within the array impacted outcome in HPTG vascular assembly studies. To assess the extent to which such a position effect exists in these data, we examined the ‘leftover’ variation of huCD31 not explained by mouse effect or condition effect (Extended Data Fig. 5). We found no significant evidence of a position effect in the arrays (Extended Data Fig. 5). Nonetheless, careful design and implementation of array layout, in which each mouse received a different layout, protected against the possibility of an undetected or misspecified position effect and enabled us to reliably screen a high number of conditions using HPTG *in vivo*.

Our results demonstrated that in the absence of human fibroblasts, the lowest weight % GelMA materials best supported robust vascular self-assembly, in a manner dependent on GelMA dose (Fig. 3d,e). Conversely and interestingly, when human fibroblasts were included in the microtissue, a larger range of material conditions (~2 – 10 wt% GelMA) supported robust vascular self-assembly, with lesser GelMA dose-dependence (Fig. 3d,e). We observed similar results for networks with host (mouse) red blood cells (TER119+; Fig. 3e). Thus, these combinatorial studies that incorporated both cellular and material variables revealed a phenomenon of stromal ‘rescue’ of higher wt% microenvironments, in which the presence of fibroblasts enabled more microenvironments to robustly support vascular self-assembly.

### Engineering designer liver tissues

We finally investigated the utility of using HPTG to identify formulations of engineered tissues that could be used as therapeutic candidates for human disease and injury. We sought to identify materials that optimally support the survival and function of primary hepatocytes encased within engineered tissues, which is an important step toward creating functional implantable tissues for treating liver disease^23^.

We first transduced hepatocytes with a lentivirus in which luciferase is expressed under the albumin promoter, to later facilitate non-destructive detection of albumin promoter activity, a surrogate for hepatic function. Transduced hepatocytes were then aggregated along with fibroblasts in microwells to create hepatic aggregates, which supports hepatic phenotype *in vitro* and survival *in vivo*^17,23,66^. Finally, we encapsulated hepatic aggregates within six different material formulations in distinct microtissue wells in the array (Fig. 4a).

**Figure 4.**
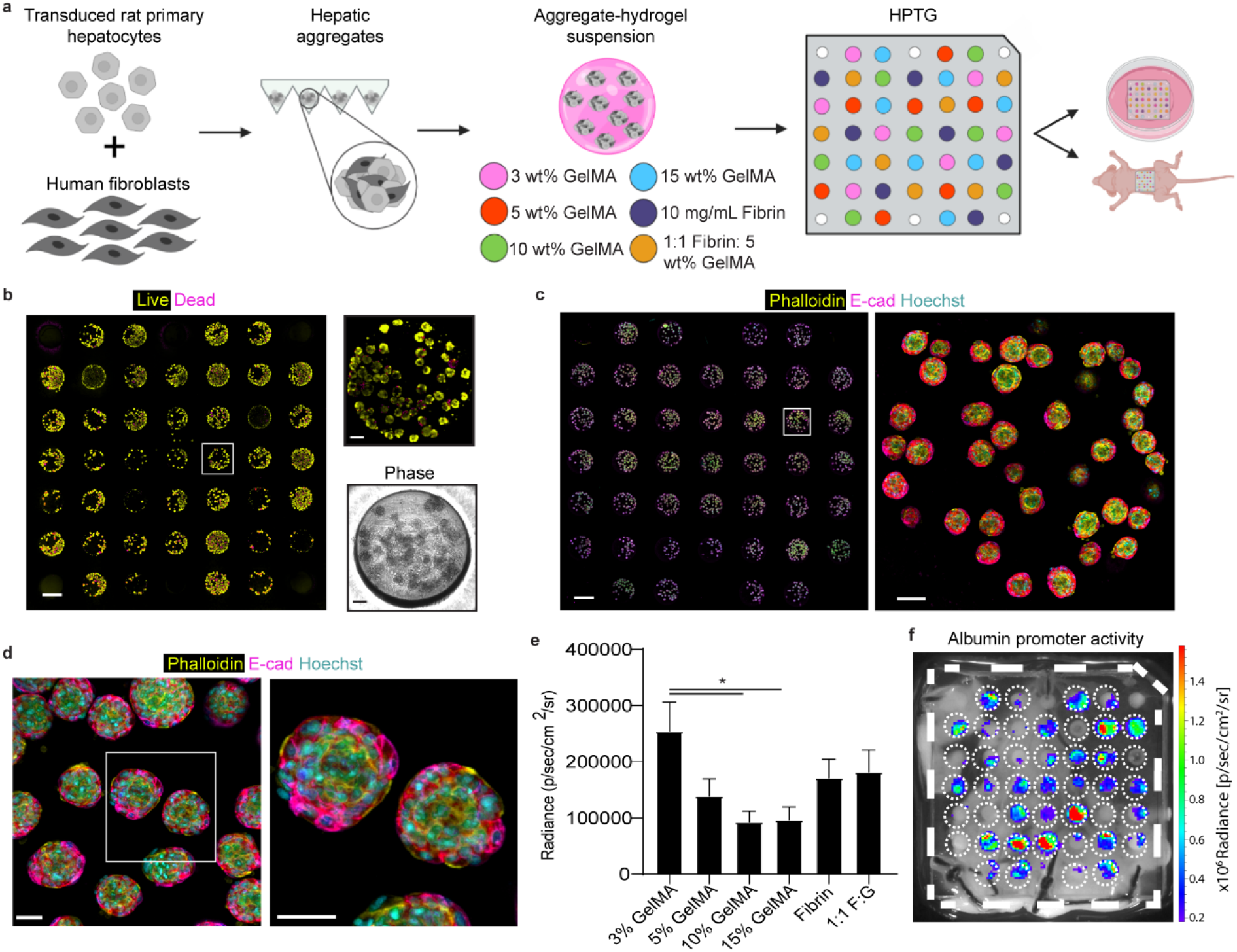
HPTG for generating and screening ‘designer’ tissues across microenvironments. Hepatocytes were aggregated with human fibroblasts to create hepatic aggregates, and aggregates were suspended in bulk in various material microenvironments (3 wt%, 5 wt%, 10 wt%, 15 wt% GelMA, 10 mg/mL fibrin, and 1:1 fibrin:5 wt% GelMA). **a,** Schematic of the experimental design. In each array there are six materials screened, each with seven technical replicates, in n=5 mouse hosts. **b,** MIP of an array filled with primary hepatocyte/NHDF aggregates (left) stained with a calcein for live (yellow) and ethidium homodimer for dead (magenta) assay. Scale bar, 1000 μm. (Right) Magnified phase image of a single well. Scale bars, 100 μm. **c,** MIP of (left) an array filled with primary hepatocyte/NHDF aggregates immunostained for E-cadherin (E-cad, magenta, epithelial cell tight junctions), phalloidin (yellow, cell actin cytoskeleton), and Hoechst (cyan), marking cell nuclei. Scale bar, 1000 μm. (Right) Magnified MIP of a single microtissue in a 5 wt% GelMA microenvironment. Scale bar, 100 μm. **d,** (Left) MIP of magnified 3D hepatic aggregates suspended in 5 wt% GelMA microenvironment within a microwell. Scale bar, 50 μm. (Right) 3D reconstruction of hepatic aggregates. Scale bar, 50 μm. **e,** Bioluminescent signal in various microenvironments after eight days in vivo. Error bars indicate SEM. *P < 0.05 by one-way analysis of variance (ANOVA) followed by Tukey’s post-hoc test. **f,** Image of albumin-driven bioluminescence expression within a representative explanted HPTG array after 8 days in vivo.

After *in vitro* encapsulation in the array, hepatic aggregates remained viable and expressed markers normally found on hepatocytes following photopolymerization within the HPTG array after 24 hours of culture (Fig. 4b,c). 3D reconstruction of the immunofluorescent signal from individual wells stained with E-cadherin and phalloidin demonstrated that hepatic aggregates retained a spherical morphology, with cells expressing E-cadherin-positive epithelial tight junctions generally localized to the perimeter of hepatic aggregates within array wells (Fig. 4c,d).

After eight days of implantation in athymic mice, we explanted the HPTG arrays and performed bioluminescence imaging to measure albumin promoter activity. We observed parallelized bioluminescent signal within individual wells across the array. In particular, the 3 wt% GelMA microenvironment supported the greatest albumin promoter activity in functioning hepatocytes after implantation, at a level twice that of the least supportive material (10 wt% GelMA, Fig. 4e,f). Thus, HPTG can be leveraged to identify tissue formulations that support the engraftment and survival of functional engineered tissues *in vivo*.

## Discussion

We report a platform for ‘plug and play’ material-and cell-based screening *in vivo*, which we call Highly Parallel Tissue Grafting (HPTG). We used 3D printing technology^17^ to print biocompatible hydrogel “well-plates”, in which a single hydrogel slab contains 43 individually addressable wells, enabling the creation of designer microtissues in which tissue components can be rapidly modified, added, or removed. HPTG enabled us to examine vascular self-assembly *in vivo*, by analyzing 688 distinct observations and 21 different material/cellular formulations using only 8 animal subjects. These studies revealed the critically important role of combinatorial effects between different microenvironmental variables in material- and cell-based studies upon engraftment *in vivo*. This entire study, including all analyses, was completed in ~1-2 months. Using previous generation technologies, this study would have been logistically infeasible.

Using HPTG, we identified photoprintable microenvironments that promote vascular self-assembly, host-graft integration, and tissue function *in vivo*. We found that different materials with different wt% supported differential self-assembly of patent vasculature. HPTG further enabled us to identify a combinatorial relationship between the inclusion of stromal cells in engineered tissues and the material that these cells are encased in. Thus, HPTG enables combinatorial *in vivo* screening studies that were not previously possible in order to identify interactions between materials and the cells they encase, providing a route towards accelerating pre-clinical safety and efficacy studies. These studies also set the stage for future mechanistic studies that could further tease apart the driving variables in the role of material formulation on vascular assembly, such as stiffness of the material, ligand density, or porosity.

Taken together, these studies demonstrate the potential power of HPTG for diverse applications, such as basic research to study vascularization, which could have important implications for fields beyond tissue engineering such as cancer biology, as well as to vascularize therapeutic tissue implants for treating disease. HPTG screening technology could also open paths toward a vast number of other medical applications, such as personalized pharmacologic screening and patient-specific therapeutic disease treatments, in areas such as heart, kidney and neurological disease.

---

**Extended Data Figure 1.**
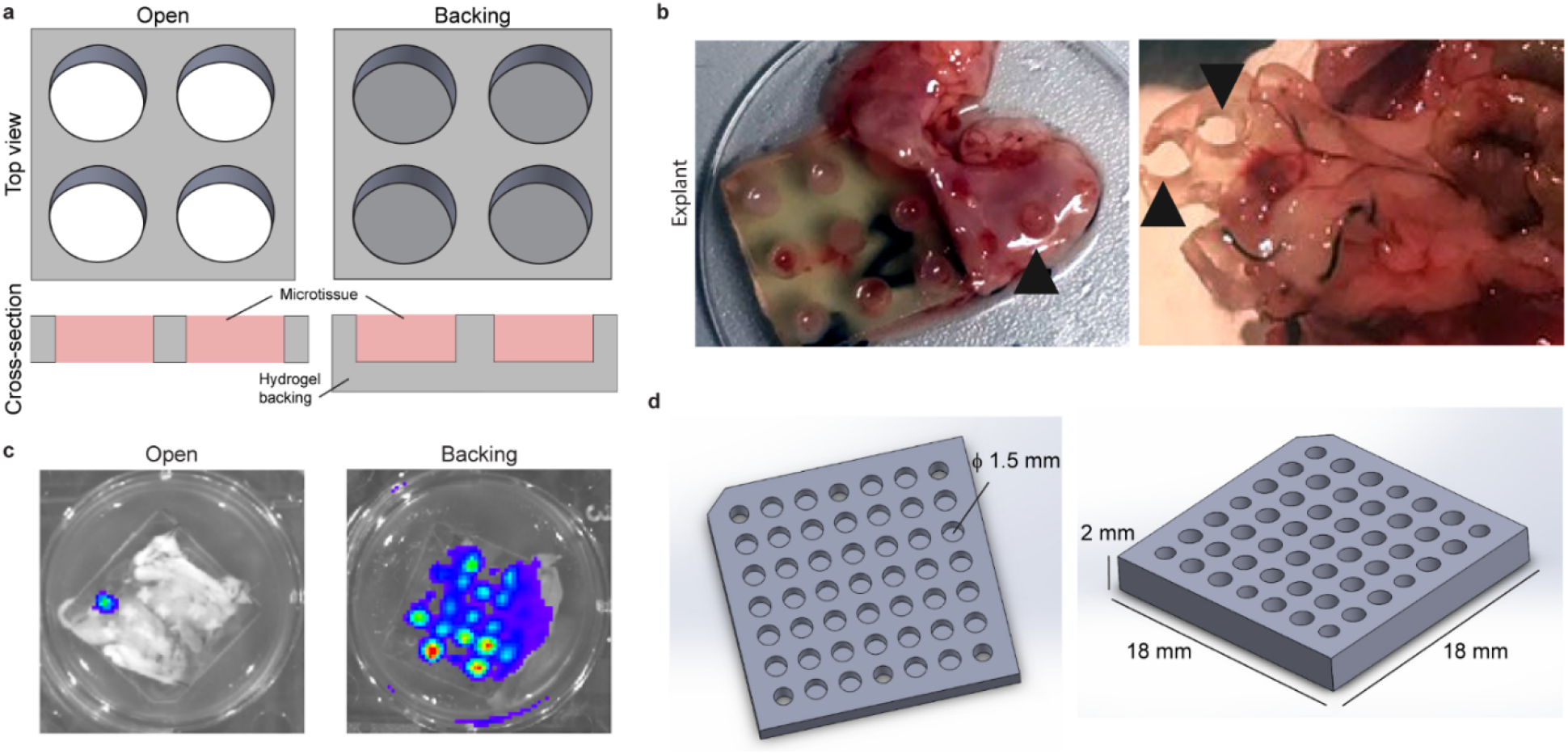
Prototyping early iterations of HPTG device for in vivo screening. **a,** (Left) Schematic of well architecture within an HPTG device with “open” wells (no backing component) and (right) with a backing component. **b,** Gross images of device architecture without a backing component directly after explant at 1 week, showing mouse tissue protruding through the array wells. Black arrows point to removal of or missing microtissue (HEK 293T cells engineered with a constitutive luciferase reporter suspended in a fibrin hydrogel) from array wells. **c,** Representative bioluminescence images of explanted HPTG arrays with open wells (left) versus those with a backing component (right). All wells of all arrays had been filled with HEK 293T cells expressing luciferase in fibrin hydrogel before implantation. **d,** Computer-aided designs of the final iteration of scaled-up HPTG device architecture with backing, 43 wells, and 6 suture holes.

**Extended Data Figure 2.**
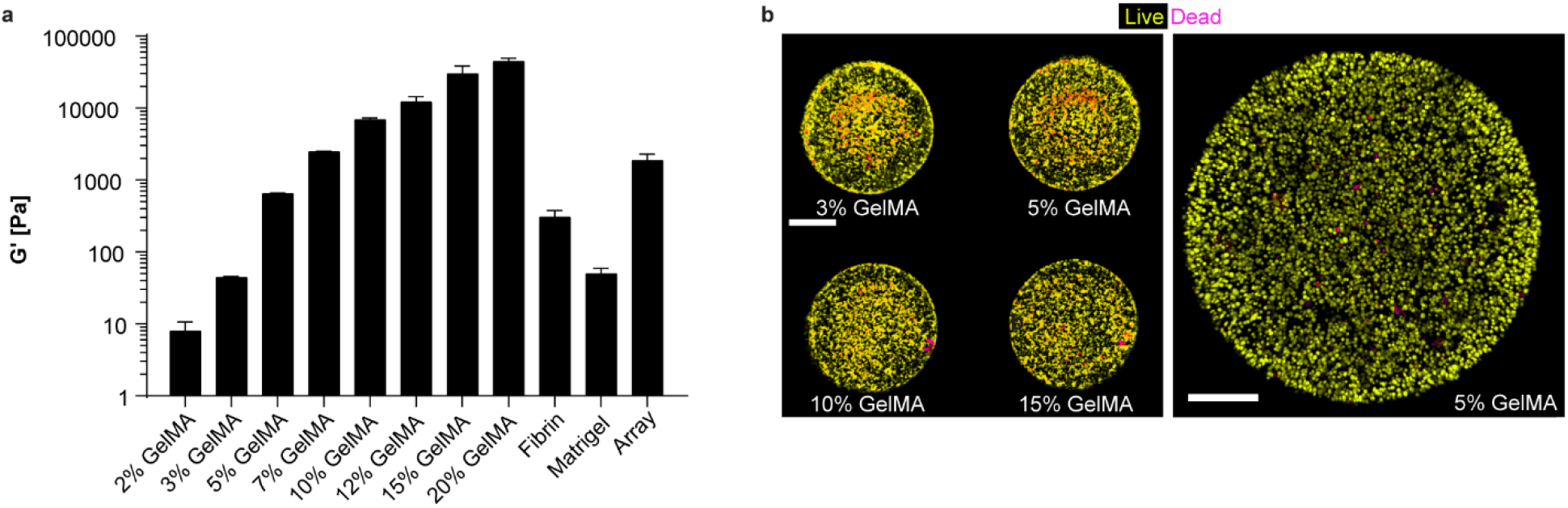
Characterization of material bioinks. **a,** Storage moduli (G’) for each of the GelMA, Fibrin, and Matrigel materials infilled into the HPTG array, as well as of the PEGDA/GelMA HPTG array scaffold, as determined via *in situ* parallel-plate rheometry. Error bars indicate SEM. **b,** HPTG array wells seeded with Human Umbilical Vein Endothelial Cells (HUVECs) in 3-15 wt% GelMA and stained for live (calcein, yellow) and dead (ethidium homodimer, magenta) cells. Scale bars, 400 μm (left) and 200 μm (right).

**Extended Data Figure 3.**
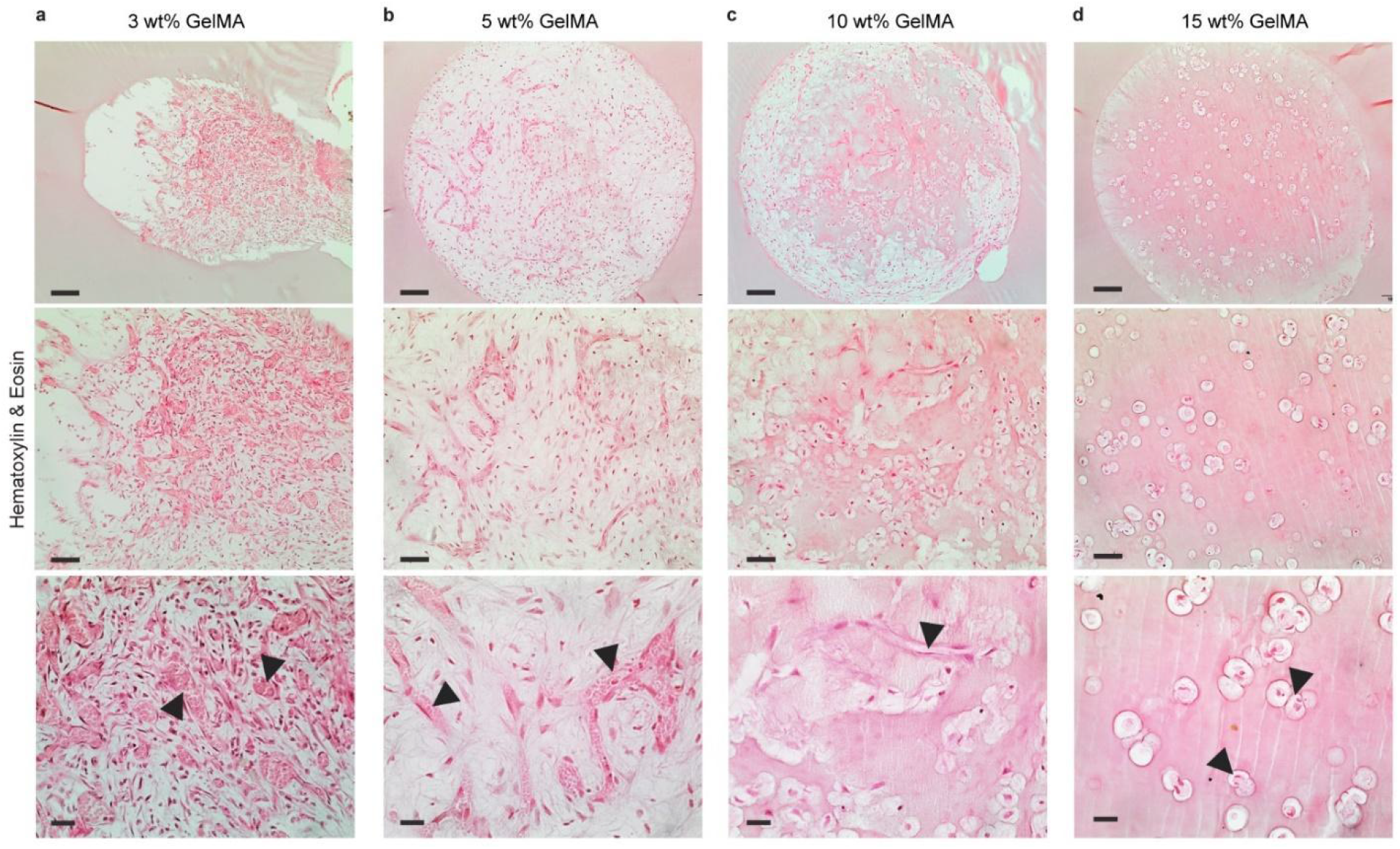
2D analysis of HPTG array. Hematoxylin and eosin (H&E) staining of 2D paraffin sections of HPTG arrays explanted after one week, screening HUVECs in **a,** 3 wt% (arrows indicate vessels filled with red blood cells; three vertical panels from top to bottom show increasing magnifications), **b,** 5 wt% (arrows indicate vessels filled with red blood cells), **c,** 10 wt% (arrow indicates hollow lumens), and **d,** 15 wt% GelMA (arrows indicate punctate endothelial cells). Scale bars top to bottom are 100 μm, 50 μm, and 20 μm.

**Extended Data Figure 4.**
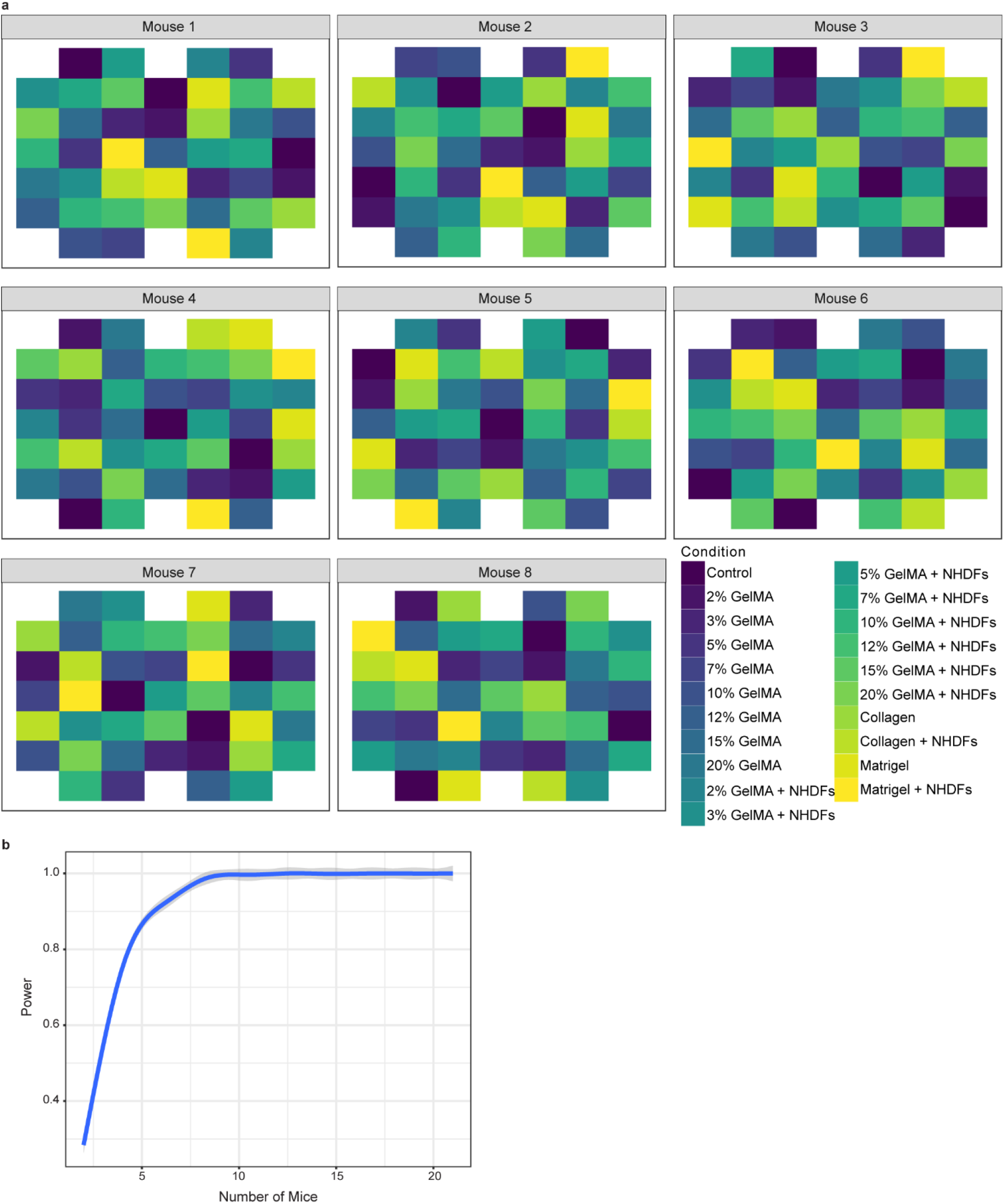
Experimental design maps for 8 HPTG arrays screening 21 conditions. **a,** The eight HPTG device well layouts is designed such that each mouse receives a different layout to protect against “position effect”, i.e., the possibility that some positions in the mouse lead to greater vascularization than others. In the first array, conditions were randomly organized across the array with the constraint that two replicates of the same condition cannot appear within two rows of each other, and two replicates of the same condition cannot be an equal number of columns from the center column. Each subsequent array layout is then a rotated version of the first one, shifted by a random number of rows and columns. **b,** For the 21-condition array design, we display the power to detect conditions that differ from the control condition, using effect sizes estimated from our preliminary data (Fig. 2).

**Extended Data Figure 5.**
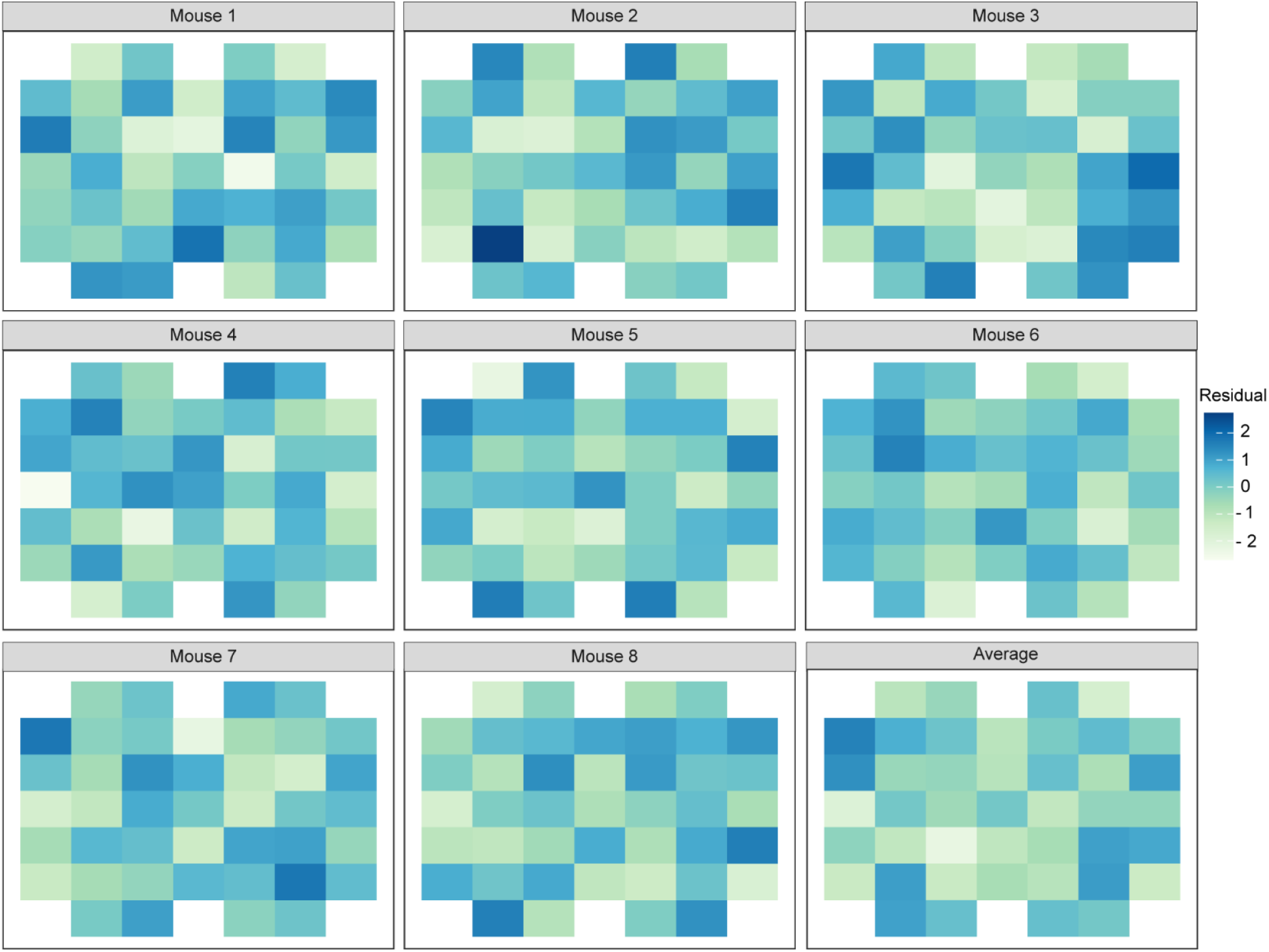
Investigating presence of location bias in next-gen HPTG design. Residual plots of huCD31+volume for each individual array across eight mice, as well as averaged across all mice (lower right), after fitting a model to predict huCD31+ volume using fixed effects for both mouse and condition. Each colored cell represents a well location on the array.

## Methods and Materials

### Design of HPTG array architectures

Computer-aided design of HPTG array architectures was designed on SolidWorks (Dassault Systems SolidWorks Corp.).

### Polymer and photoinitiator synthesis

Poly(ethylene glycol) diacrylate (PEGDA, 3.4 kDa) was prepared as previously described^67^. Lithium phenyl-2,4,6-trimethyl-benzoyl phosphinate (LAP) was synthesized as described previously^68^.

Gelatin methacrylate (GelMA) was synthesized as previously described^55^ with minor modifications^17^. Briefly, gelatin was dissolved in carbonate-bicarbonate buffer at 50°C and methacrylic anhydride was added dropwise. After 3 hours the solution was precipitated with ethanol. The precipitate was allowed to dry, dissolved in phosphate-buffered saline (PBS, Fisher), and frozen at −80°C. The GelMA was then lyophilized and stored at −20°C until use.

### Material microenvironment synthesis and rheology characterization

GelMA infill materials were formulated to contain desired wt% GelMA (2, 3, 5, 7, 10, 12, 15, 20 wt%), 10 mM LAP, and PBS. Fibrin was prepared at 10 mg/mL from Fibrinogen (Bovine Plasma, Sigma) and Thrombin (Human plasma, Sigma). Type I collagen (Fisher) was prepared at 2 mg/mL. To investigate the mechanical properties of the material microenvironments screened in the HPTG array, we ran rheological measurements on all materials in bulk on an Anton Paar MCR-301 instrument equipped with a parallel-plate geometry (diameter = 8mm) at a gap height of 500 μm. First, we determined the proper frequency and amplitude (strain) for the materials to ensure we were in the linear viscoelastic range. We found that 5% strain and 0.5 Hz fell in that range for all samples tested. We then ran 6-minute time sweeps for all the conditions from 2-20 wt% GelMA and 10 mg/mL fibrin in triplicate. We pipetted 25 μL of prepolymer directly onto the rheometer platform. The prepolymer was then equilibrated for 60 seconds, followed by a 30-second exposure of 405 nm light (Mightex BioLED, 24.5 mW/cm^2^) from the bottom of the custom-made, translucent plate to crosslink the gels. The storage (G’) and loss (G”) moduli were allowed to stabilize with the remaining time, and the last thirty seconds of readings were averaged for the final storage modulus of the sample. All trials were conducted at room temperature.

### Cell Culture

Primary human umbilical vein endothelial cells (HUVECs; Lonza; passages 4 to 7) were cultured on dishes in Endothelial Cell Growth Medium-2 (EGM-2; Lonza). Normal human dermal fibroblasts (NHDFs; Lonza; passages 4 to 10) were cultured on dishes in Dulbecco’s Modified Eagle’s Medium (DMEM; Corning) with 10% (v/v) fetal bovine serum (FBS; Gibco) and 1% (v/v) penicillin-streptomycin (pen-strep; Invitrogen).

### Rat Hepatocyte Isolation and Culture

Primary rat hepatocytes were isolated as previously described^69–72^. The hepatocytes were maintained in high-glucose DMEM (Corning) containing 10% (v/v) FBS (Gibco), 1% (v/v) insulin, transferrin, sodium selenite supplement (ITS; BD Biosciences), 7 ng/mL glucagon (Sigma), 0.04 ug/mL dexamethasone (Sigma), and 1% (v/v) pen-strep (Invitrogen). Once isolated, primary rat hepatocytes were plated in AggreWell Micromolds (400μm square AggreWell micromolds, Stem Cell Technologies) along with NHDFs at a 1:1.6 ratio and incubated in a hepatocyte medium containing DMEM with L. Glutamine (Corning), dexamethasone, FBS, ITS, and glucagon overnight at 37°C to form hepatic aggregates^23^.

### Fabrication of HPTG arrays

HPTG hydrogel arrays are printed using a custom-designed stereolithography apparatus for tissue engineering (SLATE) previously described^17^. HPTG arrays are printed under DLP light intensity of 24.5 mW cm^-2^ using a mixture of 3.25:10 wt% 3.4 kDa PEGDA: GelMA, 17 mM LAP, and 1.519 mM tartrazine (Sigma) at 50 μm layer thickness. HPTG arrays measure 18 mm X 18 mm X 2 mm. Each well is 1 mm deep and 1.5 mm in diameter. HPTG arrays are then washed to remove any unreacted monomer/tartrazine and allowed to swell to equilibrium for 3 days in PBS before further use.

### Filling of HPTG arrays

Materials screened in the HPTG array construct included any combination of Matrigel (Corning), collagen (Type I rat tail, Fisher), fibrin (10 mg/mL), 2, 3, 5, 7, 10, 12, 15, 20 wt% GelMA mixtures, or a hybrid material mixture of 1:1 10 mg/mL fibrin:5 wt% GelMA. For vascularization screening arrays, HUVECs and NHDFs were first washed with PBS to remove animal serum, detached with 0.25% trypsin solution (Corning), and spun into pellets corresponding to the number of conditions to be screened. For hepatic screening, hepatic aggregates were collected and spun into pellets corresponding to the number of conditions to be screened. All cell pellets were then resuspended into appropriate material. Cell density was calculated at 12,000 HUVECs/μL, 4,300-12,000 NHDFs/μL, and 2,700 hepatocytes/μL.

To prepare HPTG arrays for cell seeding, wells within the array were carefully aspirated of PBS. The array was then positioned in an empty dish, and each well was seeded according to the “maps” created from the computational modeling and simulations (Extended Data Fig. 4). Each well received 1 μL of material/cellular formulation. After seeding a full array, the HPTG array was placed on the SLATE and allowed to photocrosslink the photoreactive (GelMA) moieties under near-UV (405 nm) light for 30 seconds. Constructs containing fibrin, Matrigel, or collagen moieties were allowed to additionally incubate at 37°C for 30 minutes before *in vitro* use or *in vivo* implantation.

### Cell viability within HPTG array

We tested the viability of both HUVECs and hepatic aggregates (primary rat hepatocytes and NHDFs) following seeding and photocrosslinking under 405 nm light for 30 seconds. Following light exposure, we incubated the cell-laden HPTG array hydrogels with LIVE/DEAD Viability/Cytotoxicity kit reagents (Invitrogen) according to manufacturer’s instructions. Fluorescence imaging was performed on a confocal laser scanning microscope (Leica TCS SP8) in the Garvey Imaging Lab in the Institute for Stem Cells and Regenerative Medicine (ISCRM) at the University of Washington.

### *In vivo* implantation of HPTG arrays

All surgical procedures were conducted according to protocols approved by the University of Washington Animal Care and Use Committee. Male and female NCr nude mice aged 8-12 weeks old (Taconic) were anesthetized using isoflurane. HPTG array tissue constructs were sutured to the perigonadal fat pad and positioned so that the open wells faced down so that the open side of the array wells were in direct apposition to the perigonadal fat pad. The beveled corner of the array was positioned in the top lefthand corner. The incisions were closed aseptically, and the animals were administered slow releasing buprenorphine (72 hour) 1 mg/kg after surgery.

### HPTG array harvesting, processing, and immunohistochemistry

Mice were sacrificed at the termination of the experiment (7-14 days). Array constructs were harvested from the intraperitoneal space along with the engrafted perigonadal fat pad. The tissue constructs were immediately fixed following excision in 4% (v/v) paraformaldehyde (PFA; VWR) for 72 hours at 4°C and then washed with PBS for 3, 30-minute durations.

To immunostain and visualize proteins of interest in 3D, thick tissue, we used an adapted version of the Clearing Enhanced 3D (C_e_3D) method^73^. First, the excised array constructs were blocked whole overnight at room temperature in C_e_3D alternative block buffer containing 1% (w/v) bovine serum albumin (BSA, Sigma), 1% (v/v) normal donkey serum (NDS; VWR), 0.1 M Tris (Sigma), and 0.3% (v/v) Triton X-100 (Sigma) with gentle shaking. The following day the tissues were incubated in primary antibody diluted 1:100 in fresh block buffer and 5% (v/v) dimethyl sulfoxide (DMSO; Sigma; a penetration enhancer) at 37°C for 24 hours. Samples were washed for 6 hours in fresh block buffer and incubated in species-appropriate, fluorophore-conjugated secondary antibody diluted 1:500 in fresh block buffer and 5% (v/v) DMSO overnight at 37°C with gentle shaking. Phalloidin (Fisher) was added 1:100 at this step to visualize actin cytoskeleton staining. Following this incubation, samples were washed for six hours at room temperature in a wash buffer containing 0.2% (v/v) Triton X-100 and 0.5% (v/v) 1-thioglycerol (Sigma) in PBS with gentle shaking.

Immediately following immunostaining, we incubated samples in Ce3D clearing solution containing 22% (v/v) N-methylacetamide (Sigma), 80% (w/v) Histodenz (Sigma), 0.1% (v/v) Triton X-100 in PBS at room temperature for 24 hours with gentle shaking. Hoechst 33342 (Invitrogen) was added 1:500 to the C_e_3D solution to counter-stain for nuclei. The arrays were then transferred to a larger volume of fresh C_e_3D clearing solution the following day for long-term, light-protected storage at room temperature to enhance clearing. The cleared array tissue constructs were placed on glass-bottom dishes and 3D imaged overnight on a confocal laser scanning microscope (Leica TCS SP8). Image z-stacks were acquired through Leica Application Suite X (LAS X) software (Leica Microsystems). To image an optically cleared HPTG array in its entirety, the 10x objective was used, and tiles of approximately 11×11 image stacks were stitched together using the mosaic merge function within LASX.

Images were converted to Imaris image format (.ims) using Imaris File Converter 9.3.1 (Oxford Instruments) and visualized within Imaris 9.3.1 (Oxford Instruments) in 3D View or Slice Mode. Automatic Surface creation mode was used to reconstruct hepatic aggregates in 3D.

Image z-stacks of immunostained HPTG arrays were reconstructed in 3D using Vesselucida 360 software. Vesselucida 360 was then used to generate parameterized data describing the architecture of the networks (volume, surface area, length). All analyses were performed blinded using Vesselucida Explorer software and then mapped back to the screened condition according to array maps.

### HTPG array map design

To engineer an array design that protects against “position effect” (i.e., the possibility that certain regions of the array promote better vascularization), we created a different array layout for each mouse. Each of the eight arrays shown in Extended Data Fig. 4 was designed such that each condition contained two technical replicates; furthermore, there were three technical replicates of the control. The arrays were designed so that the two replicates for a given condition were not within two rows of each other and were not the same number of columns away from the center column. Moreover, each subsequent array is a rotated version of the first one, shifted by a random number of rows and columns. Based on these conditions, colored array maps were generated using statistical computing software, R.

### Linear models to account for mouse and condition

We fit linear models to estimate the effect of each condition on vessel volume measured with CD31, while including a fixed effect for each mouse to control for potential biological variation between mice. Due to the highly skewed distribution of vessel volume, we fit this model on a log scale.

Let *vol_ijk_* denote the vessel volume (huCD31) for a well in mouse i that received condition j and is located in position k. Let 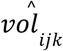 be the predicted percent vessel for a well in mouse i with condition j in position k. We first fit a model of the form

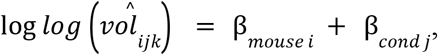

where we refer to β_*mouse i*_ as the mouse effect and β_cond j_ as the condition effect. To obtain the p-values displayed Figure 3(e), we tested the null hypothesis that the fixed effect for a given experimental condition without NHDFs equals the fixed effect for the same condition with NHDFs added. In this section, we focus on vessel volume measured with CD31, but similar models were fit for each of the other dependent variables displayed in Figure 3(e).

To obtain the eight residual maps shown in Extended Data Fig. 5, we computed the values of log 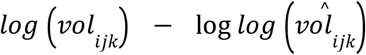 for every position in every mouse. The ‘Average’ residual map in Extended Data Fig. 5 shows the values of log 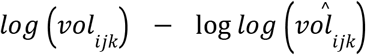 for each position averaged across mice. These residuals show leftover variation in log vessel volume after accounting for mouse and condition. Any consistent pattern across mice in these residual plots would provide evidence of a position effect. Extended Data Fig. 5 does not clearly provide such evidence.

Despite the lack of clear visual evidence of a position effect (Extended Data Fig. 5), we conducted a formal statistical test for the presence of a position effect. Our model imposed left-right symmetry on the coefficients corresponding to the positions. Furthermore, we modeled the position effect as monotone from the top to the bottom of the array. The model is as follows:

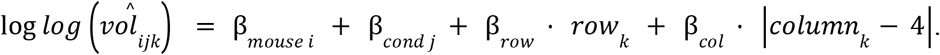

In this model, *row_k_* denotes the row corresponding to position *k* and *col_k_* denotes the column corresponding to position *k* (each range from 1 to 7). When we fit this model, neither *β_row_* nor *β_column_* were found to be statistically significantly different from 0.

### Power analysis

For each number of mice ranging from 2-21, we generated 1000 synthetic datasets. Each synthetic dataset uses arrays designed according to the layout in Extended Data Fig. 4a. Let *pva_ijk_* denote the percent vessel area for a well in mouse i with condition j in position k. The synthetic datasets are generated from the following model:

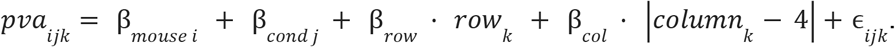

The magnitudes of the row effect, column distance effect, mouse effects, and noise (*ϵ_ijk_*) match that of the initial HPTG data (Fig. 2). In the synthetic datasets, fourteen conditions have no effect, meaning that their average percent vessel area matches that of the control condition.

Seven of the conditions (the active conditions) have an average vessel volume that is 25 percentage points higher than the control condition. This assumed effect size is smaller than the difference between 5 wt% GelMA and 15 wt% GelMA microenvironments in the initial HPTG data (Fig. 2).

After generating each synthetic data set, we fit a linear model that accounts for mouse, row effect and column distance effect, and condition effect. The estimated power is the proportion of times, across all datasets and across all active conditions, that we observe a statistically significant effect at alpha=0.05 with a Bonferroni correction for multiple comparisons. The estimated power, as a function of the number of mice, is shown in Extended Data Fig. 4b. This simulation study shows that we can achieve greater than 90% power to detect a 25 percentage point difference in vessel volume using as few as eight mice.

### Bioluminescent imaging within HPTG array

To visualize albumin-driven luciferase expression as an indirect metric for hepatic function, primary rat hepatocytes were transduced with a lentiviral vector expressing firefly luciferase under the albumin promoter (pTRIP.Alb.IVSb.IRES. tagRFP-DEST, provided through a Materials Transfer Agreement with Charles Rice, The Rockefeller University) as previously described^74^. The concentrated virus was diluted 1:5 in hepatocyte medium containing N-2-hydroxyethylpiperazine-N-2-ethane sulfonic acid buffer (HEPES; 20 mM; Gibco) and 4 μm/mL polybrene (Sigma) in ultra-low attachments 6-well plates (Corning) for 6 hours. After incubation, the transduced hepatocytes were collected for aggregating into hepatic aggregates and subsequent screening in HPTG arrays. After 1 week of implantation, arrays were explanted from the mouse host and immediately incubated with cell culture media containing D-luciferin (0.15 mg/mL; PerkinElmer) for 10 minutes and then imaged using the In Vivo Imaging System (IVIS) Spectrum imaging system (PerkinElmer) and Living Image software (PerkinElmer).

### 2D tissue histology

To process arrays for traditional 2D histology, previously fixed and tissue-cleared constructs were re-hydrated in PBS for 24 hours. The arrays were then embedded in paraffin for immunohistochemical analysis. Arrays were sectioned in 5 μm slices using a microtome and transferred to slides. For gross visualization of tissues within wells, sections were stained with hematoxylin and eosin (H&E).

## Acknowledgements

This research was supported by the NIH R01DK128551 (K.R.S.), Wellcome Leap as part of the HOPE program (K.R.S.) W.M. Keck Foundation (K.R.S.), Allen Distinguished Investigator Award (K.R.S.), a Paul G. Allen Frontiers Group advised grant of the Paul G. Allen Family Foundation (K.R.S.), NIH Maximizing Investigators’ Research Award (R35GM138036, C.A.D.), NSF CAREER Award (DMR 1652141, C.A.D.), NSF Graduate Research Fellowships (C.E.O., N.E.G., I.K.), a Ford Foundation Predoctoral Fellowship (C.E.O.), NCATS Translational Research Training Program TL1 TR002318 (C.L.F.), NIGMS Molecular Medicine Training Grant T32 GM095421 (C.L.F), and a Simons Investigator Award in Mathematical Modeling of Living Systems (D.W.) Materials used in this research (PEGDA and LAP) were generously gifted by Dr. Jordan Miller at Rice University. We also thank Dr. Miller for feedback on this work.

## References

1. We acknowledge that papers authored by women and scholars from historically excluded racial and ethnic groups are systematically under-cited. So that we are not further perpetuating this problem, we have made every attempt to reference relevant papers in a manner that is equitable in terms of gender and racial representation.

1. Mitrousis, N., Fokina, A. & Shoichet, M. S. Biomaterials for cell transplantation. Nat. Rev. Mater. 3, 441–456 (2018).

2. Li, S. et al. Hydrogels with precisely controlled integrin activation dictate vascular patterning and permeability. Nat. Mater. 16, 953–961 (2017).

3. Richardson, T. P., Peters, M. C., Ennett, A. B. & Mooney, D. J. Polymeric system for dual growth factor delivery. Nat. Biotechnol. 19, 1029–1034 (2001).

4. Shen, Y. H., Shoichet, M. S. & Radisic, M. Vascular endothelial growth factor immobilized in collagen scaffold promotes penetration and proliferation of endothelial cells. Acta Biomater. 4, 477–489 (2008).

5. Ballios, B. G. et al. A Hyaluronan-Based Injectable Hydrogel Improves the Survival and Integration of Stem Cell Progeny following Transplantation. Stem Cell Reports 4, 1031–1045 (2015).

6. Aguado, B. A., Mulyasasmita, W., Su, J., Lampe, K. J. & Heilshorn, S. C. Improving viability of stem cells during syringe needle flow through the design of hydrogel cell carriers. Tissue Eng. - Part A 18, 806–815 (2012).

7. Engler, A. J., Sen, S., Sweeney, H. L. & Discher, D. E. Matrix Elasticity Directs Stem Cell Lineage Specification. Cell 126, 677–689 (2006).

8. Park, S. H. et al. BMP2-modified injectable hydrogel for osteogenic differentiation of human periodontal ligament stem cells. Sci. Rep. 7, 1–15 (2017).

9. Moore, E. M. et al. Biomaterials direct functional B cell response in a material-specific manner. Sci. Adv. 7, 5830 (2021).

10. Jha, A. & Moore, E. Collagen-derived peptide, DGEA, inhibits pro-inflammatory macrophages in biofunctional hydrogels. J. Mater. Res. 37, 77–87 (2022).

11. Azarin, S. M. et al. In vivo capture and label-free detection of early metastatic cells. Nat. Commun. 6, 1–9 (2015).

12. Chen, T. T. et al. Anchorage of VEGF to the extracellular matrix conveys differential signaling responses to endothelial cells. J. Cell Biol. 188, 595–609 (2010).

13. Fan, V. H. et al. Tethered Epidermal Growth Factor Provides a Survival Advantage to Mesenchymal Stem Cells. Stem Cells 25, 1241–1251 (2007).

14. Gaharwar, A. K., Singh, I. & Khademhosseini, A. Engineered biomaterials for in situ tissue regeneration. Nat. Rev. Mater. 5, 686–705 (2020).

15. Bianco, P. & Robey, P. G. Stem cells in tissue engineering. Nature 414, 118–121 (2001).

16. Atala, A., Kurtis Kasper, F. & Mikos, A. G. Engineering complex tissues. Sci. Transl. Med. 4, (2012).

17. Grigoryan, B. et al. Multivascular networks and functional intravascular topologies within biocompatible hydrogels. Science (80-.). 364, 458–464 (2019).

18. Engler, A. J., Humbert, P. O., Wehrle-Haller, B. & Weaver, V. M. Multiscale Modeling of Form and Function. Science (80-.). 324, 208–212 (2009).

19. Kolesky, D. B., Homan, K. A., Skylar-Scott, M. A. & Lewis, J. A. Three-dimensional bioprinting of thick vascularized tissues. Proc. Natl. Acad. Sci. U. S. A. 113, 3179–3184 (2016).

20. Lancaster, M. A. et al. Cerebral organoids model human brain development and microcephaly. Nat. 2013 5017467 501, 373–379 (2013).

21. Takasato, M. et al. Kidney organoids from human iPS cells contain multiple lineages and model human nephrogenesis. Nature 526, 564–568 (2015).

22. Kaully, T., Kaufman-Francis, K., Lesman, A. & Levenberg, S. Vascularization—The Conduit to Viable Engineered Tissues. https://home.liebertpub.com/teb 15, 159–169 (2009).

23. Stevens, K. R. et al. In situ expansion of engineered human liver tissue in a mouse model of chronic liver disease. Sci. Transl. Med. 9, eaah5505 (2017).

24. Murphy, W. L., Peters, M. C., Kohn, D. H. & Mooney, D. J. Sustained release of vascular endothelial growth factor from mineralized poly(lactide-co-glycolide) scaffolds for tissue engineering. Biomaterials 21, 2521–2527 (2000).

25. Freeman, I. & Cohen, S. The influence of the sequential delivery of angiogenic factors from affinity-binding alginate scaffolds on vascularization. Biomaterials 30, 2122–2131 (2009).

26. O’Connor, C., Brady, E., Zheng, Y., Moore, E. & Stevens, K. R. Engineering the multiscale complexity of vascular networks. Nat. Rev. Mater. 1–15 (2022). doi:10.1038/s41578-022-00447-8

27. Baranski, J. D. et al. Geometric control of vascular networks to enhance engineered tissue integration and function. Proc. Natl. Acad. Sci. 110, 7586–7591 (2013).

28. Redd, M. A. et al. Patterned human microvascular grafts enable rapid vascularization and increase perfusion in infarcted rat hearts. Nat. Commun. 10, 1–14 (2019).

29. Levenberg, S. et al. Engineering vascularized skeletal muscle tissue. Nat. Biotechnol. 23, 879–884 (2005).

30. von Kortzfleisch, V. T. et al. Improving reproducibility in animal research by splitting the study population into several ‘mini-experiments’. Sci. Rep. 10, 1–16 (2020).

31. Voelkl, B. et al. Reproducibility of animal research in light of biological variation. Nat. Rev. Neurosci. 21, 384–393 (2020).

32. Higuera, G. A. et al. In vivo screening of extracellular matrix components produced under multiple experimental conditions implanted in one animal. Integr. Biol. 5, 889–898 (2013).

33. Driehuis, E., Kretzschmar, K. & Clevers, H. Establishment of patient-derived cancer organoids for drug-screening applications. Nat. Protoc. 15, 3380–3409 (2020).

34. Eduati, F. et al. A microfluidics platform for combinatorial drug screening on cancer biopsies. Nat. Commun. 2018 91 9, 1–13 (2018).

35. Mathews Griner, L. A. et al. High-throughput combinatorial screening identifies drugs that cooperate with ibrutinib to kill activated B-cell-like diffuse large B-cell lymphoma cells. Proc. Natl. Acad. Sci. U. S. A. 111, 2349–2354 (2014).

36. Huang, L. et al. Ductal pancreatic cancer modeling and drug screening using human pluripotent stem cell– and patient-derived tumor organoids. Nat. Med. 2015 2111 21, 1364–1371 (2015).

37. Flaim, C. J., Chien, S. & Bhatia, S. N. An extracellular matrix microarray for probing cellular differentiation. Nat. Methods 2, 119–125 (2005).

38. Beachley, V. Z. et al. Tissue matrix arrays for high-throughput screening and systems analysis of cell function. Nat. Methods 12, 1197–1204 (2015).

39. Wang, Y. et al. Development of a Photo-Crosslinking, Biodegradable GelMA/PEGDA Hydrogel for Guided Bone Regeneration Materials. Materials (Basel). 11, (2018).

40. Brady, A.-C. et al. Proangiogenic Hydrogels Within Macroporous Scaffolds Enhance Islet Engraftment in an Extrahepatic Site. Tissue Eng. Part A 19, 2544–2552 (2013).

41. Chen, X. et al. The epididymal fat pad as a transplant site for minimal islet mass. Transplantation 84, 122–125 (2007).

42. Weaver, J. D. et al. Vasculogenic hydrogel enhances islet survival, engraftment, and function in leading extrahepatic sites. Sci. Adv. 3, (2017).

43. Gama Sosa, M. A. et al. Low-level blast exposure disrupts gliovascular and neurovascular connections and induces a chronic vascular pathology in rat brain. Acta Neuropathol. Commun. 7, 6 (2019).

44. Mansour, A. A. et al. An in vivo model of functional and vascularized human brain organoids. Nat. Biotechnol. 36, 432–441 (2018).

45. Takebe, T. et al. Vascularized and functional human liver from an iPSC-derived organ bud transplant. Nature 499, 481–484 (2013).

46. Lee, A. et al. 3D bioprinting of collagen to rebuild components of the human heart. Science (80-.). 365, 482–487 (2019).

47. Levenberg, S., Golub, J. S., Amit, M., Itskovitz-Eldor, J. & Langer, R. Endothelial cells derived from human embryonic stem cells. Proc. Natl. Acad. Sci. 99, 4391–4396 (2002).

48. Schechner, J. S. et al. In vivo formation of complex microvessels lined by human endothelial cells in an immunodeficient mouse. Proc. Natl. Acad. Sci. 97, 9191–9196 (2000).

49. Stratman, A. N., Malotte, K. M., Mahan, R. D., Davis, M. J. & Davis, G. E. Pericyte recruitment during vasculogenic tube assembly stimulates endothelial basement membrane matrix formation. Blood 114, 5091–5101 (2009).

50. Song, H.-H. G. et al. Transient Support from Fibroblasts is Sufficient to Drive Functional Vascularization in Engineered Tissues. Adv. Funct. Mater. 30, 2003777 (2020).

51. Stevens, K. R. et al. Physiological function and transplantation of scaffold-free and vascularized human cardiac muscle tissue. Proc. Natl. Acad. Sci. 106, 16568–16573 (2009).

52. Koike, N. et al. Creation of long-lasting blood vessels. Nature 428, 138–139 (2004).

53. Benton, J. A., DeForest, C. A., Vivekanandan, V. & Anseth, K. S. Photocrosslinking of Gelatin Macromers to Synthesize Porous Hydrogels That Promote Valvular Interstitial Cell Function. Tissue Eng. Part A 15, 3221–3230 (2009).

54. Zhu, M. et al. Gelatin methacryloyl and its hydrogels with an exceptional degree of controllability and batch-to-batch consistency. Sci. Rep. 9, 1–13 (2019).

55. Shirahama, H., Lee, B. H., Tan, L. P. & Cho, N.-J. Precise Tuning of Facile One-Pot Gelatin Methacryloyl (GelMA) Synthesis. Nat. Publ. Gr. (2016). doi:10.1038/srep31036

56. Klak, M. et al. Irradiation with 365 nm and 405 nm wavelength shows differences in DNA damage of swine pancreatic islets. PLoS One 15, e0235052 (2020).

57. Lawrence, K. P. et al. The UV/Visible Radiation Boundary Region (385–405 nm) Damages Skin Cells and Induces “dark” Cyclobutane Pyrimidine Dimers in Human Skin in vivo. Sci. Rep. 8, 1–12 (2018).

58. Li, W., Germain, R. N. & Gerner, M. Y. Multiplex, quantitative cellular analysis in large tissue volumes with clearing-enhanced 3D microscopy (Ce3D). Proc. Natl. Acad. Sci. U. S. A. 114, E7321–E7330 (2017).

59. Cheng, G. et al. Engineered blood vessel networks connect to host vasculature via wrapping-and-tapping anastomosis. Blood 118, 4740 (2011).

60. Shaikh, F. M. et al. Fibrin: a natural biodegradable scaffold in vascular tissue engineering. Cells. Tissues. Organs 188, 333–346 (2008).

61. Suuronen, E. J. et al. Tissue-Engineered Injectable Collagen-Based Matrices for Improved Cell Delivery and Vascularization of Ischemic Tissue Using CD133+ Progenitors Expanded From the Peripheral Blood. Circulation 114, (2006).

62. Mimura, T. et al. Cultured Human Corneal Endothelial Cell Transplantation with a Collagen Sheet in a Rabbit Model. Invest. Ophthalmol. Vis. Sci. 45, 2992–2997 (2004).

63. Zhang, Y. et al. Collagen-based matrices improve the delivery of transplanted circulating progenitor cells: development and demonstration by ex vivo radionuclide cell labeling and in vivo tracking with positron-emission tomography. Circ. Cardiovasc. Imaging 1, 197–204 (2008).

64. Christman, K. L. et al. Injectable Fibrin Scaffold Improves Cell Transplant Survival, Reduces Infarct Expansion, and Induces Neovasculature Formation in Ischemic Myocardium. J. Am. Coll. Cardiol. 44, 654–660 (2004).

65. Bensaïd, W. et al. A biodegradable fibrin scaffold for mesenchymal stem cell transplantation. Biomaterials 24, 2497–2502 (2003).

66. Stevens, K. R. et al. InVERT molding for scalable control of tissue microarchitecture. Nat. Commun. 4, 1–11 (2013).

67. Miller, J. S. et al. Bioactive hydrogels made from step-growth derived PEG–peptide macromers. Biomaterials 31, 3736–3743 (2010).

68. Fairbanks, B. D., Schwartz, M. P., Bowman, C. N. & Anseth, K. S. Photoinitiated polymerization of PEG-diacrylate with lithium phenyl-2,4,6-trimethylbenzoylphosphinate: polymerization rate and cytocompatibility. Biomaterials 30, 6702–6707 (2009).

69. Dunn, J. C. Y., Tompkins, R. G. & Yarmush, M. L. Long-Term in Vitro Function of Adult Hepatocytes in a Collagen Sandwich Configuration. Biotechnol. Prog. 7, 237–245 (1991).

70. Seglen, P. O. Chapter 4 Preparation of Isolated Rat Liver Cells. Methods Cell Biol. 13, 29–83 (1976).

71. Chen, A. A. et al. Humanized mice with ectopic artificial liver tissues. Proc. Natl. Acad. Sci. U. S. A. 108, 11842–11847 (2011).

72. March, S., Hui, E. E., Underhill, G. H., Khetani, S. & Bhatia, S. N. Microenvironmental regulation of the sinusoidal endothelial cell phenotype in vitro. Hepatology 50, 920–928 (2009).

73. Li, W., Germain, R. N. & Gerner, M. Y. Multiplex, quantitative cellular analysis in large tissue volumes with clearing-enhanced 3D microscopy (Ce3D). Proc. Natl. Acad. Sci. U. S. A. 114, E7321–E7330 (2017).

74. Andrus, L. et al. Expression of paramyxovirus V proteins promotes replication and spread of hepatitis C virus in cultures of primary human fetal liver cells. Hepatology 54, 1901–1912 (2011).

